# Beyond the Leaderboard: Leveraging Predictive Modeling for Protein-Ligand Insights and Discovery

**DOI:** 10.1101/2025.05.12.653449

**Authors:** Dan Kalifa, Kira Radinsky, Eric Horvitz

**Affiliations:** Technion; Microsoft

## Abstract

**Motivation:** Ligands are biomolecules that bind to specific sites on target proteins, often inducing conformational changes important in the protein’s function. Knowledge about ligand interactions with proteins is fundamental to understanding biological mechanisms and advancing drug discovery. Traditional protein language models focus on amino acid sequences and three-dimensional structures, overlooking the structural and functional changes induced by protein-ligand interactions. We investigate the value of integrating ligand-protein binding data in several predictive challenges and leverage findings to frame research directions and questions.

**Results:** We show how the integration of protein-ligand interaction data in protein representation learning can increase predictive power. We evaluate the methodology across diverse biological tasks, demonstrating consistent improvements over state-of-the-art models. We further demonstrate how the study of the specific boosts in predictive capabilities coming with the introduction of the ligand modality can serve to focus attention and provide insights on biological mechanisms. By leveraging large pretrained protein language models and enriching them with interaction-specific features through a tailored learning process, we capture functional and structural nuances of proteins in their biochemical context.

**Availability and implementation:** The full code and data are freely available at https://github.com/kalifadan/ProtLigand.

## 1 INTRODUCTION

Proteins rarely act in isolation. Whether catalysing metabolic reactions, propagating signals, or regulating gene expression, most proteins rely on small-molecule ligands—cofactors, metabolites, metal ions—to assume the conformations and physicochemical states that make biological activity possible. Yet the protein language models (PLMs) that now underpin much of structural bioinformatics are trained on amino-acid sequences [18] or AlphaFold-derived coordinates [27], disregarding the ligands that stabilise folds, switch functional states, and determine specificity. As PLMs developed for protein structure, interaction, and design lack information about ligands, they share a set of blind spots: they cannot distinguish paralogues that share more than 80% sequence identity yet bind different metabolites [16]. Thus, they fail to explain why an identical fold can be thermostable in one organism but unstable in another; and they struggle to predict which mutations will dramatically change drug affinity. In short, omitting ligand chemistry is a source of deficits in today’s models when it comes to connecting sequence and structure with the functional logic of living systems.

We first introduce *ProtLigand*, a novel general-purpose PLM that learns a protein in the chemical context of the ligand it binds. Then, we show via a set of case studies how we can harness gains in predictive power to frame research directions and garner insights about specific biochemical phenomena, including cofactor-driven stability shifts and ligand-mediated conformational selection.

During pre-training, ProtLigand receives three synchronous inputs: the amino-acid sequence, a coarse backbone trace from AlphaFold2 [15] and a Simplified Molecular Input Line Entry System (SMILES) [35] string that encodes the cognate ligand. The model is trained to recover masked residues while jointly attending to the ligand, so the resulting representation is shaped not only by evolutionary and geometric constraints but also by the electrostatic, hydrophobic, and steric properties imposed by binding partners. Full architectural details—tokenization, attention scheme, and optimization objective—are provided in the Methods section. Here, we note that the approach demands no docking coordinates, only sequence, predicted backbone, and ligand identity.

Protein–ligand binding is almost always modeled through docking pipelines in which separate neural encoders are trained for the protein and the small molecule, and a downstream module then predicts the pose. Uni-Mol [42], BindNet [9], GNINA [19] and DiffDock [5] all follow this two-stream recipe, so the protein representation itself is learned without ever “seeing” its ligand. To date, no method has allowed ligand chemistry to shape the protein embedding during language-model pre-training. The only partial step in that direction, SMILESVec [23], merely averages fingerprints of known binders and therefore loses all protein sequence- and structure-specific nuance characteristic of the protein itself. Its dependence on comprehensive ligand annotations—an assumption rarely satisfied in practice, particularly in tasks such as protein–protein interaction prediction—limits its practical applicability and generalizability.

Evaluated on six benchmarks, including thermostability prediction, human protein–protein interaction classification, and metal ion binding prediction, ProtLigand outperforms the strongest sequence–structure baseline SaProt [27] across all tasks with a statistically significant improvement (Table 1). We note that numerical gains in performance coincide with biologically coherent patterns. Heme oxygenase 2 and p38 kinases, whose activities depend on porphyrin cofactors, are correctly classified only when ligand information is present. Autophagy partners ATG7 and ATG10 are recognised as interacting, whereas unrelated nuclear proteins CBX4 and BOLL are correctly rejected—cases in which SaProt fails, apparently misled by structural similarity. Removing ligand features produces a larger accuracy drop on the thermostability set than ablating any purely structural feature, implying that ligand chemistry likely plays a causal role in thermal behaviour. We show how we can harness the predictive model to be a hypothesis generator: hypotheses and research directions can be framed by examining the selective sensitivity of predictive performance on specific ligand signals. Interactions that are highly sensitive to the introduction of ligand information are promising candidates for experimental validation.

**Table 1:** Experimental results on six downstream tasks. Statistically significant results with *p <* 0.05 using a two-tailed paired t-test (with Holm–Bonferroni correction) ^1^ are marked with an asterisk (*). For ProtLigand, we report the mean and standard deviation over 3 independent fine-tuning runs (with different seeds) ^2^. The best result is highlighted in bold.

| Model | HumanPPI | Thermostability | Metal Ion Binding | DeepLoc (Acc%) |  | EC |
| --- | --- | --- | --- | --- | --- | --- |
| | (Acc%) | (Spearman’s $\rho$) | (Acc%) | Binary | Subcellular | ($F_{max}$) |
| MIF-ST [38] | 75.54 | 0.694 | 75.08 | 91.76 | 78.96 | 0.807 |
| ESM-1b [24] | 82.22 | 0.708 | 73.57 | 92.83 | 80.33 | 0.864 |
| ESM-2 [18] | 76.67 | 0.680 | 71.56 | 91.96 | 82.09 | 0.868 |
| GearNet [40] | 73.86 | 0.571 | 71.26 | 89.18 | 69.45 | 0.874 |
| ESM-GearNet [39] | 84.09 | 0.651 | 74.11 | 92.94 | 82.30 | 0.882 |
| SaProt [27] | 86.67 | 0.710 | 75.75 | 93.55 | 83.19 | 0.876 |
| **ProtLigand** | **90.00\*±0.5** | **0.731\*±0.02** | **77.54\*±0.18** | **94.02\*±0.21** | **83.88\*±0.07** | **0.887\*±0.002** |

<sup>1</sup> Significance markers refer to Bonferroni-adjusted p-values. Full statistics are reported in Supplementary Table 3.
<sup>2</sup> 95% confidence intervals for ProtLigand are provided in Supplementary Table 4.

ProtLigand also can be employed to explore the “dark matter” of unknown protein-ligand interactions. We introduce a novel generator, trained separately from the PLM, that has the ability to extrapolate a plausible ligand fingerprint from the protein representation and decodes that encoded description back to a SMILES string. Figure 1 illustrates this capacity: given only the protein, the model proposes ligands held out from the training set whose functional groups match those of experimentally verified binders. In summary, ProtLigand fuses sequence, structure, and ligand context into a single representation that improves prediction accuracy, yields interpretable biochemical insights, and extends naturally to orphan targets. By embedding the chemistry of ligand binding partners directly into protein language models, we enrich the representations of protein chemistry to more faithfully reflect the molecular logic of life.

**Figure 1.**
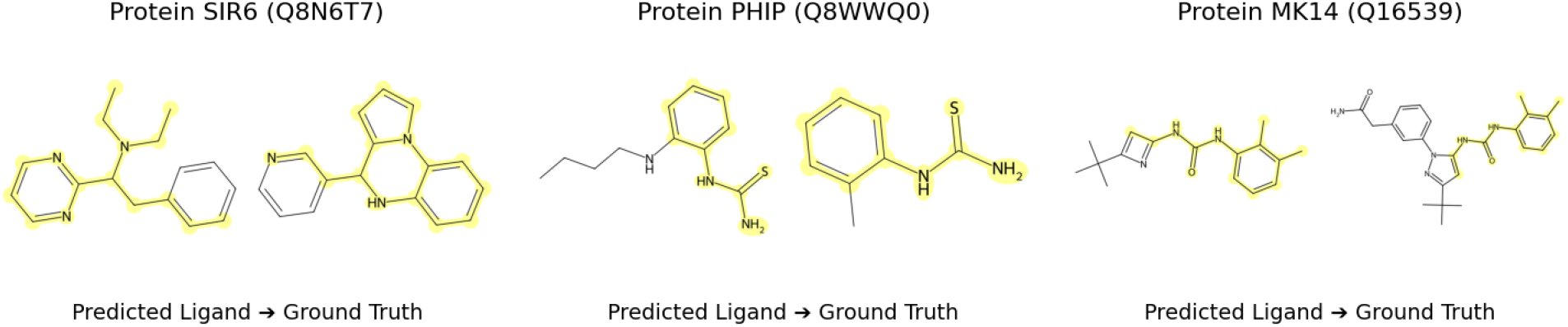
Examples of ligands successfully predicted for various unseen proteins. Given only a protein as input, the ligand generator produces a proxy representation of the predicted interacting ligand. The ligand decoder then predicts the SMILES strings to obtain the chemical structures of these ligands. Highlighted regions show functional groups that match between the generated ligands and the ground truth labels.

The contributions of this work are threefold: (1) We introduce ProtLigand, the first general-purpose protein language model that incorporates ligand context via a cross-attention mechanism, yielding representations that capture not only sequence and structure but also the biochemical effects of binding partners, and show state-of-the-art predictive performance. (2) We demonstrate a methodology for extending the model’s utility from pure predictions to providing proposals for de novo ligands. (3) We show how the considerations of sensitivity of predictive power to ligand information can provide hypotheses about mechanisms: We achieve new top results on a suite of six downstream tasks—including thermostability prediction, HumanPPI classification, and metal-ion binding. Additionally, we show case studies that link performance gains with ligand information to specific biochemical phenomena.

## 2 MATERIALS AND METHODS

### 2.1 Protein-Ligand Dataset

ProtLigand is pre-trained on the PDBbind (v.2020, CC BY 4.0) dataset [34], a widely used dataset for protein–ligand interaction modeling [5, 9, 19, 42]. It contains experimentally validated protein–ligand complexes, categorized by protein types. We construct protein–ligand pairs by linking each protein to its corresponding interacting ligand from the dataset.

To ensure reliable performance during training, we partition the dataset into fixed training and validation sets using the Graph-Part algorithm [30], which performs homology-aware sequence clustering. Sequences in the validation set share no more than 30% Needleman–Wunsch sequence identity with any sequence in the training set, enabling a strict assessment of generalization. The reported model performance is evaluated on benchmark test sets that are strictly held out and share less than 30% Needleman–Wunsch sequence identity with any sequence in the pre-training dataset (including both training and validation sets). This ensures that all evaluation sequences are effectively unseen during inference, supporting a fair and rigorous evaluation. Following the procedure used in SaProt [27], we retrieve AlphaFold2 structures [15] for each protein from AlphaFoldDB [32] using their UniProt IDs. Low-confidence regions (with pLDDT scores below 70) in AlphaFold2-predicted structures were removed using the same filtering strategy described in SaProt [27]. Overall, this process results in a complete dataset of proteins comprising sequence and structure and their corresponding interacting ligands. The final dataset comprises 17,393 protein–ligand pairs (14,331 for training and 3,062 for validation), covering 3,347 unique proteins across 12 protein families. See Supplementary Section D.1 for an overview of protein categories and associated ligand types in the dataset, highlighting the chemical diversity of these protein–ligand complexes.

### 2.2 Overview of ProtLigand

#### ProtLigand

is a novel ligand-aware PLM (See Figure 2) that enhances protein representations by leveraging information from interacting ligands. The model is trained on amino acid sequences along with their 3D structures. We construct a structure-aware sequence by combining residue and structural tokens and apply masked language modeling (MLM) [8]. A pre-trained base PLM (SaProt 650M AF2 version [27]) generates a contextual representation for the masked sequence, while the known interacting ligand is passed through a ligand encoder (ChemBERTa-77M-MLM [3]) to obtain representations, which are linearly projected into the protein’s latent space. To capture the biochemical context of proteinligand interactions, ProtLigand incorporates a cross-attention mechanism [33] between the protein and ligand representations. Through this attention, ligand-derived keys and values inform protein sequence queries, enabling a refined protein representation that reflects ligand-binding effects. The refined representation is used to reconstruct the masked residues through a language model head. See Supplementary Material A for a detailed description of the algorithm, along with implementation specifics and computational complexity.

**Figure 2.**
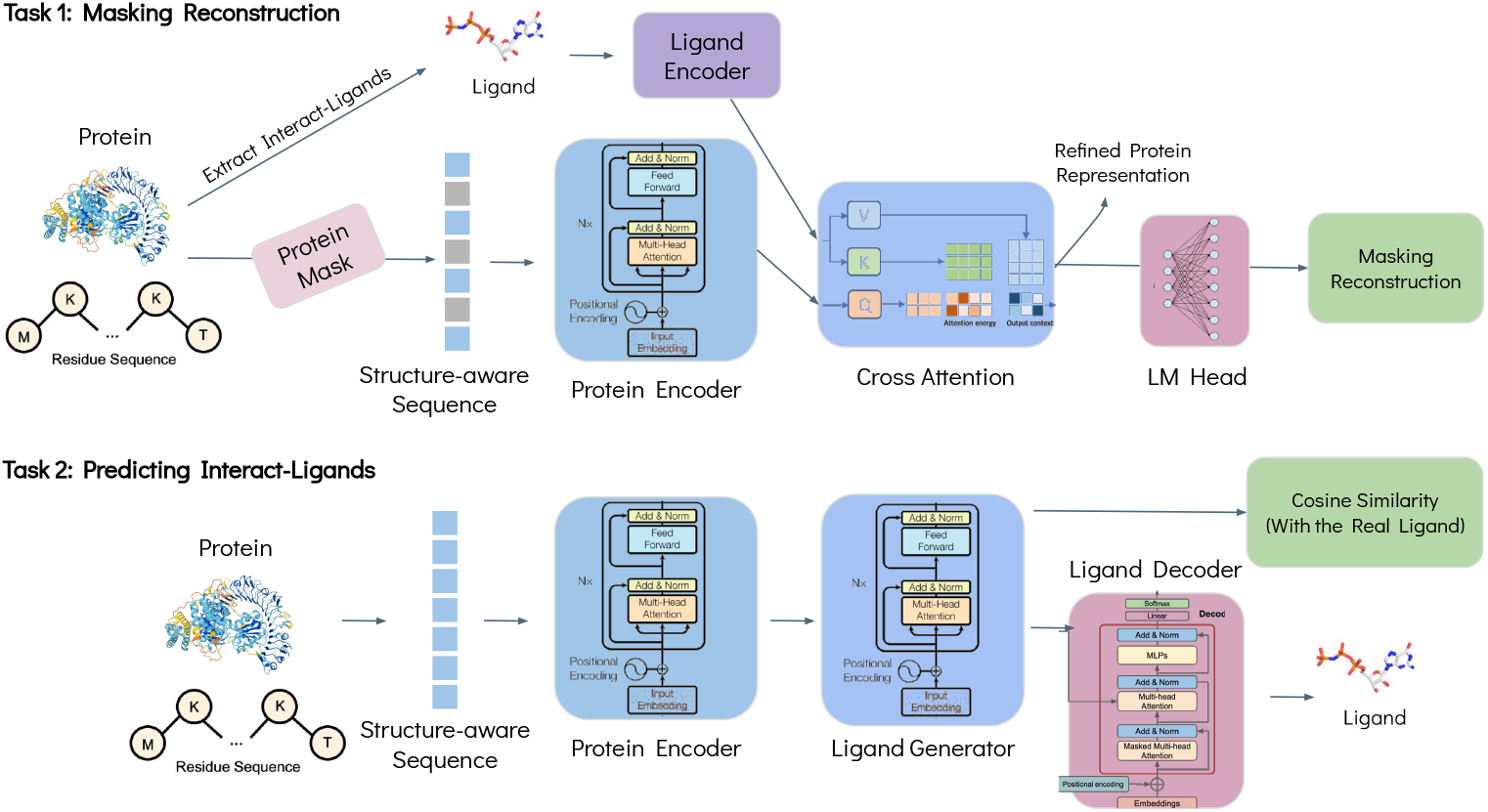
ProtLigand pre-training architecture. The model is trained on amino acid sequences along with their 3D structures, retrieved using AlphaFold2. We construct a structure-aware sequence by combining residue and structural tokens and apply masked language modeling. The protein encoder generates a contextual representation for the masked sequence. In parallel, the known interacting ligand is passed through a ligand encoder to obtain its representation, which is linearly projected into the protein’s latent space. A cross-attention mechanism refines the protein representation using ligand-derived keys and values. The refined representation is used to reconstruct the masked residues via an LM head. The protein representation can be harnessed as a lightweight novel ligand generator, trained alongside the PLM, that extrapolates a plausible ligand fingerprint from the protein representation and decodes that fingerprint back to a SMILES string. This approach allows the model to learn ligand awareness even in cases where ligand information may be missing at inference time.

### 2.3 Ligand Generator

We extend the applicability of ProtLigand to scenarios where ligand information is unavailable by introducing a novel **ligand generator**, trained separately from the PLM (See Supplementary Material A.3 for details). This component predicts a proxy representation of a protein’s interacting ligand based solely on its internal representation, using cosine similarity between the predicted and true ligand representations as the training objective. Then, during inference, we can utilize this generator to learn representations for unseen proteins. Such a generation ability can help to identify expected structures of ligands that have not yet been identified and of potential synthetic ligands that might be generated as candidate agonists and antagonists to known protein-ligand interactions.

To better understand the representations that are generated, we incorporate a **ligand decoder** (See Supplementary Material A.4 for details), which reconstructs the original chemical ligand sequence (encoded in the SMILES representation) from the learned ligand latent space. This enables the model to propose candidate ligands for unseen proteins, potentially uncovering novel interactions. To evaluate the generated ligands, after training the decoder on the training set of the Protein-Ligands dataset, we generate interacting ligands for proteins from the unseen set and compare the generated ligands to the ground-truth interacting ligands (See Figure 1).

### 2.4 Benchmark Tasks

We evaluate our proposed method on a comprehensive set of downstream tasks, selected according to the latest SOTA benchmark used by SaProt [27], to assess the effectiveness of our approach for protein representation learning. These tasks span several biological fields, from protein-protein interaction prediction to thermostability estimation and binding site classification. To ensure the validity of the results, we validate all downstream task splits by enforcing that sequences in one set share no more than 30% Needleman–Wunsch sequence identity with any sequence in the other sets. This homology filtering ensures minimal overlap between training and evaluation data, minimizing the risk of data leakage.

#### Protein-Protein Interaction Prediction

Reliable detection of protein–protein interactions (PPIs) is essential for deciphering cellular processes and uncovering therapeutic targets, especially when the interactions have not been previously characterized [14]. We use the HumanPPI dataset from the PEER benchmark [37] to evaluate binary interaction prediction between protein pairs. Model performance is assessed using classification accuracy.

#### Protein Function Prediction

We assess protein function using two tasks: thermostability prediction on the “Human-cell” split of the FLIP benchmark [7], formulated as a regression task and evaluated using Spearman’s rank correlation coefficient (*ρ*); and Metal Ion Binding [13], a binary classification task designed to predict the presence of metal ion–binding sites within a protein, evaluated by accuracy.

#### Protein Localization Prediction

We adopt the DeepLoc dataset [1], which provides two variants of the subcellular localization task: (1) a binary classification task and (2) a 10-class multiclass classification task. We use accuracy as the primary performance metric for both variants.

#### Protein Annotation Prediction

We evaluate protein functional annotation using the Enzyme Commission (EC) number prediction task from the DeepFRI benchmark [11]. This task is formulated as a multi-label classification problem, where each protein may be assigned one or more EC labels. *F*_max_ score is used for evaluation.

## 3 RESULTS

We present ProtLigand’s empirical performance across a diverse set of benchmark tasks, demonstrating SOTA results.^1^ In addition to quantitative metrics, we provide qualitative analyses that link performance gains to specific biochemical mechanisms. Comprehensive ablation studies, included in Supplementary Material C, evaluate the contribution of each architectural component to the overall model performance, including a sequence identity analysis to assess generalization beyond the pre-training distribution.

We report the mean and standard deviation of ProtLigand’s performance over three independent fine-tuning runs (with different random seeds) to ensure reproducibility (Table 1). Additionally, we validate the statistical significance of performance differences, using a two-tailed paired t-test with a 95% confidence level (*p <* 0.05), comparing observations from the tested models. The normality of the paired differences was confirmed using the Shapiro–Wilk test [25]. Supplementary Table 8 lists raw p-values for each of three runs and the Holm–Bonferroni value applied to the family of three; the starred metric in Table 1 reflects this adjusted p.

### 3.1 Benchmarks Results

In Table 1, we compare the performance of ProtLigand with six SOTA baseline methods (See Supplementary Section A.6 for details) across six diverse biological downstream tasks. In particular, ProtLigand significantly outperforms all baseline methods, achieving the highest score across all tasks. We calculate the Cohen’s d [4] effect sizes to quantify the magnitude of ProtLigand’s improvements on each task, and observe large effect sizes (*d >* 0.8) across the board. We attribute the boosts to the inclusion of the ligand modality.

The most significant performance gains are observed on the HumanPPI and Thermostability benchmarks, tasks where protein–ligand interactions are biologically central and ligand annotations are more available. On HumanPPI, ProtLigand improves accuracy from 86.67% (SaProt) to 90.00%. On the Thermostability task, it achieves a higher Spearman’s *ρ* (0.731 vs. 0.710). These results suggest that ProtLigand may better capture biological features relevant to these tasks, which are overlooked in sequence and structure models.

Smaller, yet consistent, gains are observed in Metal Ion Binding, DeepLoc, and EC prediction tasks, where ligand-binding information is sparse or not explicitly present in downstream data. Nevertheless, ProtLigand maintains an edge across all metrics, suggesting that its learned representations generalize even in the absence of direct ligand supervision.

The narrower gap on DeepLoc and EC, compared to HumanPPI or Thermostability, aligns with the nature of these tasks: while structural context is crucial, ligand-driven modulation is known to be less prominent. This further supports the view that ProtLigand is particularly effective in domains where ligand binding has been confirmed to play a regulatory or stabilizing role.

Overall, these results underscore the value of incorporating lig- and context into protein modeling. ProtLigand’s consistent gains across structurally and functionally diverse tasks highlight the value of multimodal learning and demonstrate that ligand-aware representations provide unique, generalizable signals for protein function prediction.

### 3.2 Ligand Context Boosts Prediction Confidence

The performance gap observed in downstream tasks, such as HumanPPI, reflects only the difference in success rates, by cases where ProtLigand correctly predicts interactions and SaProt does not. However, another crucial aspect highlighted in Figure 3 is ProtLigand’s consistently higher prediction *confidence* across most of its predictions. This confidence-driven distinction emphasizes the model’s ability to make more decisive and biologically meaningful predictions.

**Figure 3.**
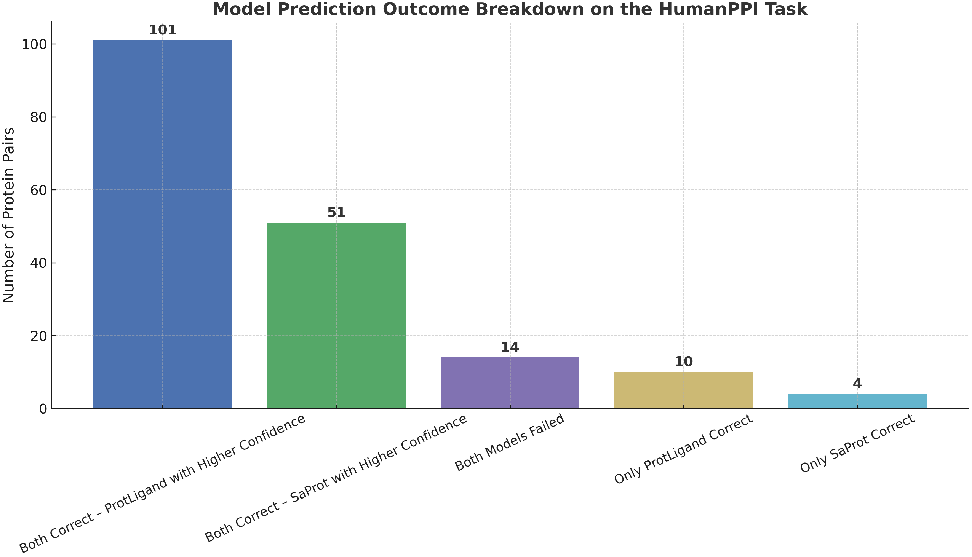
Breakdown of the ProtLigand prediction outcomes on the HumanPPI test set on a representative run with a fixed random seed for reproducibility. Each bar represents the number of protein pairs grouped by correctness and confidence: whether both models were correct, failed, or only one model predicted correctly, further split by which model was more confident.

As classification networks must not only be accurate but also reliable in their uncertainty estimates, calibrated confidence [12] is essential for interpretability and downstream decision-making. We evaluate and plot the expected calibration error (ECE) [12] for the HumanPPI task and the expected normalized calibration error (ENCE) [17] for the Thermostability regression task, using 10 equal-width bins over the confidence or predicted value range, following the calibration equation [12, 17]. ProtLigand consistently exhibits improved calibration over SaProt, with lower ECE in classification (0.043 vs. 0.070) and reduced ENCE in regression (0.048 vs. 0.276). Reliability diagrams (see Supplementary Figure 5) show that ProtLigand tracks the identity line more closely than SaProt across the confidence bins.

A key insight of our approach is that ligand information introduces additional functional context that complements traditional protein-level representations. Although prior models often capture structural and evolutionary features, the learned representations typically lack detailed information about how proteins interact with small molecules, information that is critical for many tasks, such as binding affinity prediction or drug response modeling.

Ligand signals encode important biochemical constraints, including charge distribution, hydrophobicity, and 3D conformation, all of which influence binding pocket compatibility. Incorporating these signals allows the model to better differentiate between proteins with similar sequences but divergent functions (e.g., paralogs) that cannot be distinguished through sequence alone. For properties like thermostability, ligand properties can have stabilizing or destabilizing effects on the protein–ligand complex. Including ligand features enables the model to reason about these interactions and capture their impact on protein function.

Overall, the ligand serves as a task-specific lens, enabling the model to refine its predictions based on real molecular interactions rather than relying solely on static structural features, resulting in higher confidence of the model on its predictions, leading to performance improvements as reflected in the results on the benchmarks.

### 3.3 Leaderboards to Insights

ProtLigand consistently outperforms SaProt across all downstream tasks, as shown in Table 1. When performance gains are consistently driven by biologically meaningful inputs, such as ligand-based features, this may signal that those inputs capture relationships of functional relevance. We note that we cannot take selective boosts in the predictive capabilities, marking causal interactions. It is important to avoid conflating prediction with causation; a model’s ability to predict biological outcomes more accurately does not in itself support causal inferences. On the other hand, significant boosts on specific classes of prediction can provide hints that can frame hypotheses, questions, and follow-on studies.

Ligands can be viewed as rich sources of mutual information between the protein input and target biological properties. Even if the model does not encode causal structure explicitly, selective improvements resulting from the inclusion of ligand context suggest that the additional information helps capture biochemical properties—such as charge distribution, hydrophobicity, or structural complementarity—that are associated with observed outcomes like stability or binding.

For instance, ProtLigand’s improvement on the Thermostability task suggests that ligand-derived features contain information relevant to thermal behavior. While this does not establish that ligands causally stabilize proteins, it highlights a potential functional link to investigate. The model acts as a hypothesis generator, surfacing biochemical dependencies that could inform experimental follow-up aimed at establishing causality.

More broadly, when the removal of a feature leads to a disproportionate drop in performance, it may indicate the feature’s functional relevance. We see such findings as putative causal signals: while the results may be based on spurious associations, they may nonetheless provide a basis for generating biological hypotheses. In this way, predictive modeling can serve not only as a tool for performance benchmarking but also as a guide for identifying biologically meaningful variables and prioritizing experimental validation.

#### 3.3.1 Lens on Biological Insights?

The improvements of ProtLigand over existing baselines align with clear biological patterns, particularly in proteins whose function, stability, or interactions are mediated by ligand binding. Unlike SaProt [27], which relies solely on sequence and structure, ProtLigand leverages ligand context through a cross-attention mechanism, yielding representations that better reflect real-world biophysical phenomena, which enables exploration of unique biological insights.

We now compare the predictions of ProtLigand to the latest SOTA model, SaProt [27], and present reflections on biological insights stimulated by the examination of the largest discrepancies in prediction scores between baseline models and models boosted with ligand data. We categorize all cases into distinct biological groups based on their shared characteristics. These examples can provide insights about the underlying factors contributing to ProtLigand’s improved performance.

##### Heme-binding and Ligand-modulated Enzymes

ProtLigand effectively captures regulatory behavior in metallo-proteins and enzymes influenced by heme or other cofactors. In both the Thermostability and HumanPPI tasks, heme oxygenase-2 (P30519), which plays a central role in heme catabolism and depends on heme for both activity and structural integrity [10], was correctly identified by ProtLigand, with significantly higher confidence than prior baselines. Another success in both tasks was MAP kinase p38 gamma (O43924), a signaling protein whose activation and conformation are influenced by ligand-mediated phosphorylation cascades [6]. ProtLigand’s attention-based modeling likely enhances sensitivity to such ligand-mediated modulation, particularly in cases where static structural similarity alone may be misleading.

##### Dynamic Interaction-Modulated Proteins

We see signals that ProtLigand captures ligand-dependent structural flexibility, significantly enhancing interaction predictions. In the HumanPPI task, ProtLigand correctly identified that CBX4 (O00257), a SUMO E3 ligase involved in chromatin organization and DNA damage response [20], and BOLL (Q8N9W6), an RNA-binding protein essential for meiosis in germ cells [28], do not interact. Despite both being nuclear-localized, these proteins function in distinct cellular programs—CBX4 is active in somatic cell nuclei, while BOLL is germline-specific and cytoplasmically enriched. SaProt misclassified this pair with high confidence (0.90), likely due to misleading structural or sequence-level similarity.

In contrast, ProtLigand’s ligand-informed model recognized the functional incompatibility and the lack of shared ligand-driven interaction context, assigning a low score (0.04). Conversely, ProtLigand correctly predicted a functional interaction between ATG10 (Q9H0Y0) and ATG7 (O95352), key autophagy enzymes that collaborate in ubiquitin-like conjugation systems [22, 29]. This interaction requires conformational alignment at flexible binding sites, a dynamic that SaProt missed (0.92 vs. 0.40). ProtLigand likely captures these transient conformational compatibilities by leveraging ligand-associated contextual signals. Together, these cases highlight ProtLigand’s ability to help resolve interaction-relevant structural states that depend not only on static features but also on functional context and ligand-induced modulation, enabling it to outperform conventional PLMs in both sensitivity and specificity.

##### Metabolically Coupled Proteins

ProtLigand reveals biologically meaningful interactions between proteins that participate in distinct yet metabolically interconnected processes. In the HumanPPI task, ProtLigand confidently identified an interaction between ATP synthase subunit gamma (P36542), a mitochondrial enzyme essential for ATP synthesis, and carbonic anhydrase 12 (CA12, O43570), a membrane-associated enzyme involved in pH regulation [2, 36]. While SaProt assigned a low interaction score (0.35), ProtLigand predicted this pair with high confidence (0.91), likely due to its ligand-informed representation learning, which captures functional dependencies beyond sequence similarity.

A similar pattern is observed with the large ribosomal subunit protein bL9m (Q9BYD2), a mitochondrial ribosomal component associated with translational control in metabolically active states. For couples including this protein, ProtLigand achieved significant improvements over SaProt (0.52 vs. 0.09).

Although localized in different compartments, these proteins contribute to coordinated metabolic adaptations, particularly in proliferative or stress environments where ATP production and proton transport must be tightly regulated.

These findings highlight ProtLigand’s ability to capture latent biochemical coordination between functionally coupled proteins, offering insights into pathways that are often overlooked by structure or sequence-only models.

##### Ligand-Dependent Signal and Assembly Modulators

ProtLigand captures biologically meaningful interactions involving proteins whose activity is regulated by ligand binding, either through allosteric signal transduction or ligand-facilitated complex formation.

In the HumanPPI benchmark, ProtLigand confidently predicted the interaction between parathyroid hormone-related protein (P12272) and its receptor, parathyroid hormone 1 receptor (Q03431). Ligand binding to PTH1R triggers conformational changes that initiate downstream signaling pathways involved in calcium regulation and bone metabolism [41]. While SaProt assigned a low confidence score of 0.02 to this pair, ProtLigand correctly identified the interaction with a score of 0.96.

In another example, ProtLigand detected a high-confidence interaction between COP9 signalosome complex subunit 6 (Q7L5N1) and cytochrome c oxidase copper chaperone (COX17, Q14061). COX17 plays a key role in delivering copper ions for the proper assembly of mitochondrial cytochrome c oxidase, a critical enzyme in oxidative phosphorylation. Although these proteins operate in different cellular compartments, their coordinated function in cellular stress response and mitochondrial regulation has been suggested. ProtLigand assigned a confidence of 0.98, compared to SaProt’s 0.85.

These cases illustrate ProtLigand’s strength in identifying ligand-mediated relationships that depend on conformational communication or cofactor transport, functional dependencies often missed by traditional sequence- or structure-only approaches.

### 3.4 Generating Testable Hypotheses

How might ProtLigand’s boosts in predictive capabilities over a default model trained without ligand information frame mechanistically grounded, experimentally testable hypotheses? pid-mediated thermostabilization of amphipathic surfaces in Perilipin-2. The example testable hypothesis highlights how ProtLigand’s ligand-aware modeling could help with formulating studies of the functional roles of small-molecule interactions and the execution of experiments to test the hypotheses with standard biochemical and cell-biological assays.

#### Ligand-Stabilized Thermostability of Perilipin-2

ProtLigand identifies a significant stabilizing ligand interaction with Perilipin-2 (PLIN2, Q99541), a lipid droplet–associated protein. In the Thermostability task, ProtLigand achieves high correlation with experimental data (Spearman’s *ρ* ≈ 0.85), markedly outperforming SaProt (*ρ* ≈ 0.50), suggesting that ProtLigand captures key stabilizing features not explained by sequence or structure alone.

PLIN2 is known to bind lipids through a conserved pocket in its C-terminal domain (see Figure 4). Prior work by Najt et al. [21] identified this region as key for lipid binding, but its stabilizing effect on the protein’s structure has not been fully explored. ProtLigand’s accuracy implies that lipid binding may not only be functional but also thermodynamically stabilizing, locking the protein into a more rigid and ordered conformation.

**Figure 4.**
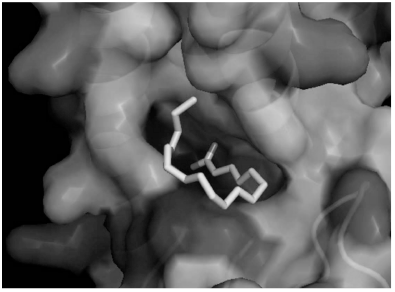
Surface rendering of the PLIN2 C-terminal domain showing the conserved lipid-binding pocket.

**Figure 5.**
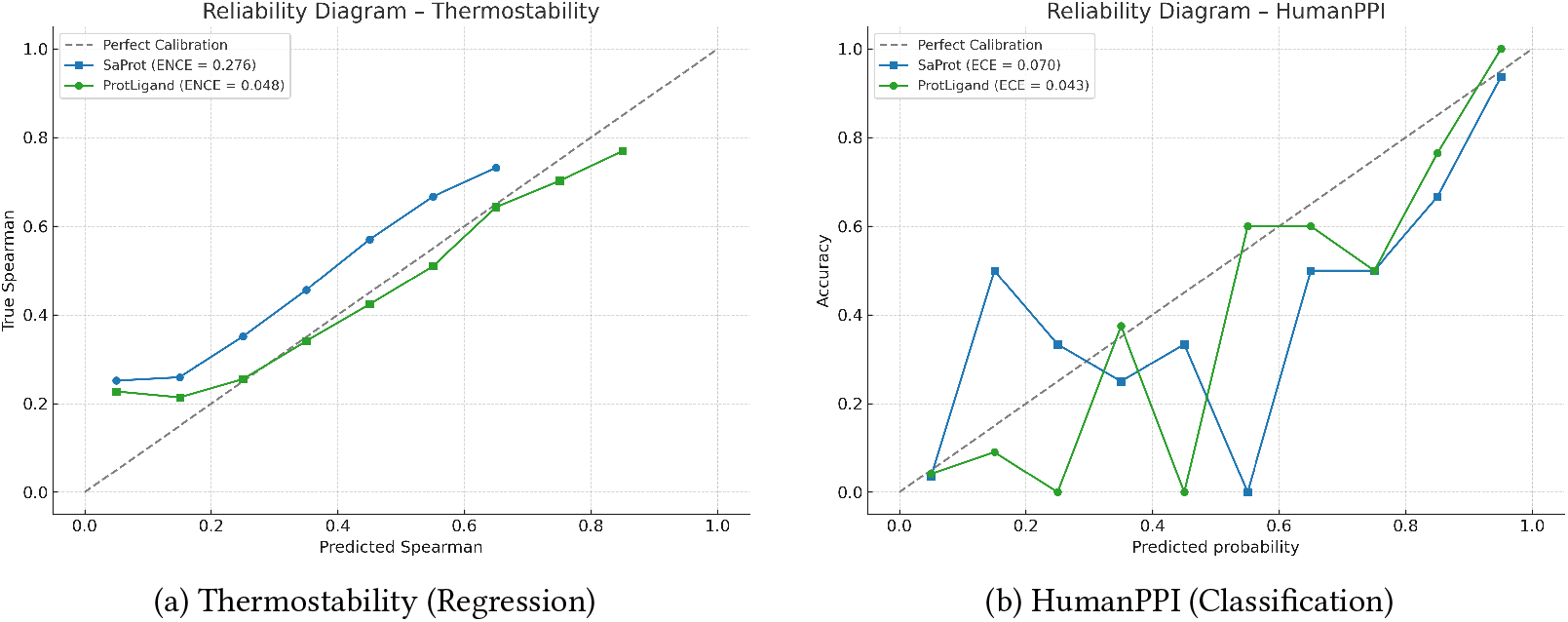
Reliability diagrams for Thermostability and HumanPPI. ProtLigand (green) demonstrates better calibration than SaProt (blue) in both tasks. In the regression task, ProtLigand achieves lower ENCE, and in the classification task, it exhibits lower ECE while closely following the perfect line.

##### Hypothesis

Binding of a lipid-like small molecule increases PLIN2 thermostability by stabilizing a flexible loop or amphipathic region in its C-terminal domain.

##### Wet-lab assay A – thermostability

Measure the thermal unfolding profile of recombinant PLIN2 in the presence versus absence of endogenous lipid ligands using differential scanning fluorimetry (DSF) or circular dichroism (CD).

##### Expected read-out

A reproducible shift in melting temperature (e.g., Δ*T*_*m*_ *>* 3^°^C) upon ligand binding, confirming increased stability.

##### Wet-lab assay B – conformational effect

Perform limited proteolysis assays with and without lipid co-incubation to detect protection patterns in predicted binding regions.

##### Expected read-out

Ligand reduces proteolytic cleavage in flexible domains, supporting a stabilization mechanism.

Lab-based experimentation would help to validate ProtLigand’s prediction and provide mechanistic insight into lipid-mediated regulation of PLIN2 structure. We hope to see the heterogeneous boosts in the predictive capabilities of models harnessed in such formulations of hypotheses, followed by experimental validation.

## 4 DISCUSSION

We introduced ProtLigand, a learning architecture and respective model, that improves PLMs by integrating ligand information during amino acid–level representation learning. The methodology enables the incorporation of both intrinsic structural properties and extrinsic interaction signals. By dynamically contextualizing proteins with their ligand environments, ProtLigand learns unified representations that capture interaction propensity, conformational flexibility, and ligand-induced stabilization, features often overlooked by conventional PLMs.

ProtLigand demonstrates strong performance in a variety of bioinformatics tasks. In particular, its improvements extend beyond specific benchmarks, indicating that the learned representations encode generalizable biophysical principles rather than task-specific heuristics. Furthermore, certain protein families, such as kinases and nuclear receptors, show particularly pronounced benefits, underscoring the value of ligand-aware modeling in biologically complex and pharmacologically relevant systems.

The presented methods and results provide a promising direction for advancing protein modeling. By bridging representations based on sequence and structure information with ligand-contextual information, the approach enables more accurate functional inference, improved interpretability, and myriad applications in structural biology and drug discovery, including the identification of new therapeutic targets and the rational design of protein–ligand interactions.

ProtLigand opens several promising directions for future research. One avenue is to leverage the generated interact-ligand SMILES representations as candidates for novel drug discovery, particularly by systematically generating ligands for PPI that currently lack known binders. Prior study has focused on the potential for attentional biases in protein science leading to blindspots with identification of interactions [26]. The use of ProtLigand as a generator can help to probe the “dark matter” of the unknowns of protein-ligand interaction.

Leveraging ProtLigand as a generator could help to expand the druggable proteome by targeting proteins that have historically been difficult to address. Another direction is to consider biology’s efficient reuse of molecules in different settings; we can explore via predictive modeling the potential role of known ligand interactions across all PPI proteins to assess whether existing compounds might align with previously unassociated proteins, potentially revealing opportunities for drug repurposing or uncovering structural similarities. We leave the systematic exploration of these strategies to future work.

### 4.1 Limitations

Although ProtLigand brings ligand awareness to protein language modeling, several limitations remain. ProtLigand is still trained on a corpus heavily enriched for lipids and drug-like scaffolds from PDBbind, so metal-containing molecules and many primary metabolites are sparsely represented. Definitive coverage across the full spectrum of cellular chemistry has yet to be demonstrated. Also, because cross-attention is applied to entire proteins rather than explicit pocket coordinates, ProtLigand cannot rank alternative binding poses or replace rigorous structure-based affinity scoring, and its inference time is approximately 21% longer compared to a sequence-only baseline. We note that, while the ligand generator produces chemically valid molecules in most cases, wetlab validation remains essential before pursuing these compounds experimentally. Finally, the cofactor-mediated stability shifts and ligand-dependent conformational selections described in our case studies have not yet been verified biochemically.

### 4.2 Ethics

The ability to generate novel small molecules raises dual-use and intellectual property concerns. To mitigate these risks, each generated structure should be passed through a three-layer safety pipeline. First, assess toxicity using tools such as Tox21, with any compound predicted to be acutely toxic, carcinogenic, or endocrine-disrupting withheld. Second, SMILES strings should be screened for detecting dangerous sequences and chemicals of concern. Third, exact-match and similarity checks should be performed against SureChEMBL and USPTO to identify previously described compounds. These measures are intended to balance open scientific dissemination with responsible stewardship of a technology that could otherwise be misused.

## A PROTLIGAND ALGORITHM

We introduce **ProtLigand**, a novel ligand-aware PLM (See Figure 2) that enhances protein representations by leveraging information from interacting ligands.

Let a protein *p* = (*S, R*) be defined by its amino acid sequence *S* = (*s*_1_, *s*_2_, …, *s*_*n*_) and its 3D structure *R*, where *s*_*i*_ *∈* 𝒱 denotes the *i*-th residue and 𝒱 is the standard residue alphabet. We then use Foldseek [31] to convert each protein structure into a sequence of 3Di tokens *F* = (*f*_1_, *f*_2_, …, *f*_*n*_) aligned with *S*, where each structural token *f*_*i*_ *∈* ℱfrom a structure alphabet ℱ. Then, we construct a structure-aware sequence *P* = (*s*_1_ *f*_1_, *s*_2_ *f*_2_, …, *s*_*n*_ *f*_*n*_), where *s*_*i*_ *f*_*i*_ ∈ 𝒱 × ℱtoken combines residue identity and local structural context. This fused sequence can then be fed into a standard Transformer encoder as basic input.

### A.1 Training Phase

During training, each structure-aware protein sequence *P* is passed through a protein encoder, producing a contextual representation *z*_*p*_ *∈* ℝ ^*N* ×*D*^, where *N* is the sequence length and *D* is the feature dimension.

We initialize the encoder with a large pre-trained PLM, SaProt-650M [27], leveraging its strong contextual understanding of structure-aware sequences. Using SaProt allows us to fairly isolate and assess the contribution of ligand integration. Alternatively, other advanced PLMs can also be used within the proposed framework.

For each protein, we retrieve its known interacting ligand from our pre-training dataset (see Section 2.1). The ligand is represented as a SMILES string *L* = (*l*_1_, *l*_2_, …, *l*_*M*_) and passed through a pretrained molecular encoder (ChemBERTa-77M-MLM [3]), producing a chemically meaningful ligand representation 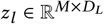, where *M* is the ligand sequence length and *D*_*L*_ is its feature dimension.

To align the ligand and protein representations in a shared latent space, ProtLigand applies an affine transformation to map the ligand representation to the protein space:

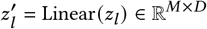

Next, we apply cross-attention [33] between the protein and lig- and tokens. The ligand-derived keys and values guide the protein’s queries, refining its representation to reflect ligand-binding context. The attention mechanism is defined as:

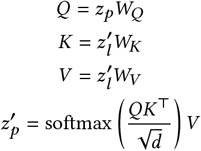

Here, *W*_*Q*_, *W*_*K*_, *W*_*V*_ *∈* ℝ^*D*×*D*^ are learnable projection matrices. The resulting 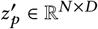 is a ligand-aware protein representation that captures the biochemical influence of ligand interactions.

### A.2 Masked Language Modeling Training Objective

For each input protein, 15% of the amino acids are randomly masked. Each masked amino acid *s*_(*i*)_ has an 80% chance of being masked for prediction, 10% chance of replacement with a random amino acid, and 10% chance of remaining unchanged. Suppose the number of masked amino acids is *N*, the training objective ℒ_*MLM*_ is to minimize:

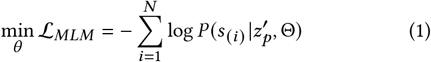

where 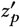 is the ligand-aware protein representation and Θ are the parameters of the ProtLigand model.

### A.3 Ligand Generator

During inference, experimentally determined ligand information is often unavailable. To eliminate this dependency, we introduce a ligand generator module trained separately during the pre-training phase. This component learns to predict a proxy representation of the interacting ligand directly from the protein representation.

The ligand generator is implemented as a Transformer encoder with *L*_1_ layers and *H*_1_ heads each, followed by a linear projection. Given the protein representation *z*_*p*_ *∈* ℝ^*N* ×*D*^ produced by the protein encoder, the generator computes:

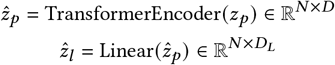

where *D*_*L*_ is the ligand representation dimension, aligned with the output of the pre-trained ligand encoder. To assess the quality of the generated ligand representations, we minimize the cosine distance between the predicted ligand representation 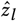 and the ground-truth ligand representation derived from the ligand encoder *z*_*l*_ :

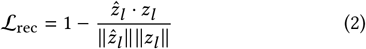

This loss encourages the protein encoder to capture ligand-relevant features, allowing ProtLigand to operate effectively even in the absence of known ligand data at inference time.

### A.4 Ligand Decoder

The ligand decoder is trained to reconstruct the SMILES sequence from the generated ligand representation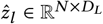. It consists of an embedding layer, learnable positional encoding, a Transformer decoder with *L*_2_ layers and *H*_2_ heads each, and a linear output layer with output dimension equal to the vocabulary size.

Given a target ligand sequence *t*, the decoder computes:

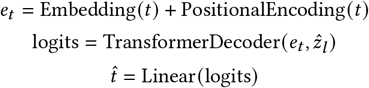

To assess the quality of the SMILES reconstruction, we minimize the cross-entropy loss between the predicted token distribution and the ground-truth SMILES sequence:

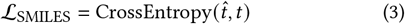

We trained the ligand decoder alongside the ligand generator, minimizing the mutual loss ℒ= ℒ _SMILES_ + ℒ _rec_.

### A.5 Inference Phase

During inference, we leverage the ligand generator (See Section A.3) to produce the interacting ligand representation from the protein representation, eliminating the need for known ligand input. Then, given the ligand representation and the protein, ProtLigand produces the ligand-aware protein representation. Furthermore, task-specific classification heads are incorporated to enable predictions for the specific downstream tasks. This allows the model to remain ligand-aware without external annotations and generalize to unseen proteins. To decode the interacting ligand representations into SMILES sequences, we generate molecules autoregressively using top-*k* sampling (*k* = 3), until a termination token is produced. The chemical validity of each generated ligand is verified using RDKit. Representative examples of valid generations are shown in Figure 1.

### A.6 Baselines

Incorporating SOTA models as baselines and following SaProt [27] for fair comparison, we include ESM-1b [24] and ESM-2 [18], which are considered top-performing sequence-based models. Structure-based baselines include GearNet [40] and MIF-ST [38], while ESM-GearNet [39] serves as a representative joint sequence–structure model. We also evaluate against SaProt itself [27], the current leading PLM that incorporates AlphaFold-derived structure tokens.

### A.7 Implementation Details

#### A.7.1 Pre-Training Phase

We utilized the pre-trained SaProt 650M [27] as the base protein encoder and ChemBERTa-77M-MLM [3] as the base ligand encoder. Following ESM-2 [18] and BERT [8], 15% of the tokens are randomly masked during training.

The hyperparameters of both ProtLigand and the ligand generator (summarized in Table 2) were selected via grid search based on performance on the validation set of the pre-training dataset. As a result, the encoder and decoder are configured with *L*_1_ = *L*_2_ = 6 layers and *H*_1_ = *H*_2_ = 8 attention heads per layer. The dimensionalities of the protein and ligand representations are set to *D* = 1280 and *D*_*L*_ = 768, respectively. To accommodate long protein sequences, inputs are truncated to a maximum of 1,024 tokens. All training is performed using mixed-precision arithmetic to improve memory efficiency and computational throughput.

**Table 2:**
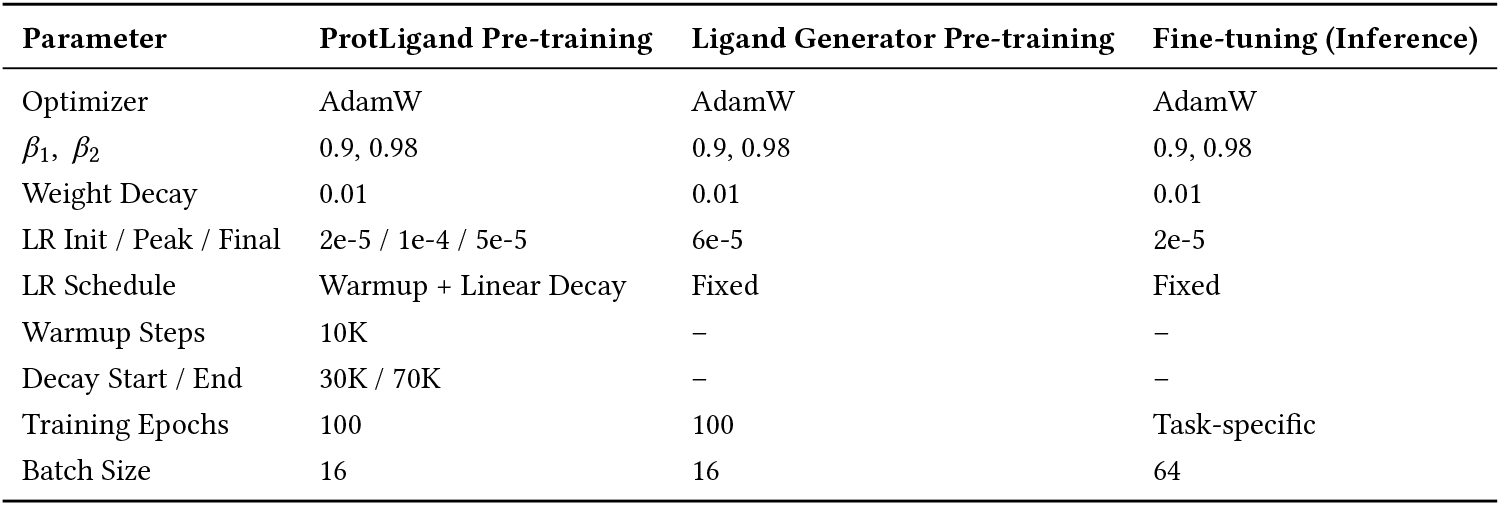
Training hyperparameters for ProtLigand and its ligand generator.

| Parameter | ProtLigand Pre-training | Ligand Generator Pre-training | Fine-tuning (Inference) |
| --- | --- | --- | --- |
| Optimizer | AdamW | AdamW | AdamW |
| $\beta_1, \beta_2$ | 0.9, 0.98 | 0.9, 0.98 | 0.9, 0.98 |
| Weight Decay | 0.01 | 0.01 | 0.01 |
| LR Init / Peak / Final | 2e-5 / 1e-4 / 5e-5 | 6e-5 | 2e-5 |
| LR Schedule | Warmup + Linear Decay | Fixed | Fixed |
| Warmup Steps | 10K | – | – |
| Decay Start / End | 30K / 70K | – | – |
| Training Epochs | 100 | 100 | Task-specific |
| Batch Size | 16 | 16 | 64 |

#### A.7.2 Fine-Tuning Phase

During inference, we incorporate task-specific classification heads to enable predictions for these tasks. Furthermore, following SaProt [27] settings, we conducted evaluations of our model and all baselines using the same set of hyper-parameters reported in SaProt to ensure fair comparisons. These hyperparameters are summarized in Table 2.

#### A.7.3 Computational Complexity

All experiments were conducted on 4× NVIDIA A100 80GB GPUs. ProtLigand’s pre-training required approximately 72 hours (288 GPU hours), with most overhead stemming from its ligand generator and cross-attention modules. In contrast, SaProt [27] was trained from scratch over three months using 64 GPUs. Thus, ProtLigand achieves a 99% reduction in pretraining cost by leveraging a pretrained encoder and integrating ligand context through lightweight modules.

At inference time, ProtLigand introduces only modest over-head. SaProt inference takes approximately 0.014 seconds per 1,000 residues, whereas ProtLigand processes the same input in about 0.017 seconds—representing a 21% increase. This overhead is minimal given the improved predictive performance and full reuse of pre-trained PLM infrastructure.

## B STATISTICAL TESTS

### B.1 Statistical Significance Tests

We perform paired two-tailed t-tests to assess the statistical significance of ProtLigand’s improvements over the top-performing baseline, SaProt [27], across all six benchmark tasks. Holm–Bonferroni correction is applied across 3 runs; the maximum adjusted p-value is reported per task. The raw and adjusted *p*-values are summarized in Table 3. Statistically significant differences (*p <* 0.05) indicate that the observed performance improvements are unlikely to have occurred due to stochastic variation.

**Table 3:** Raw and Holm–Bonferroni-adjusted *p*-values for ProtLigand and SaProt across all tasks. Raw *p*-values are computed using a paired two-tailed t-test over per-sample prediction scores across 3 fine-tuning runs.

| Task | Raw $p$-values | Adjusted $p$-value |
| --- | --- | --- |
| HumanPPI | 0.0120, 0.0140, 0.0130 | 0.0360 |
| Thermostability | 0.0003, 0.0001, 0.0002 | 0.0004 |
| Metal Ion Binding | 0.0150, 0.0170, 0.0160 | 0.0450 |
| DeepLoc Binary | 0.0090, 0.0110, 0.0100 | 0.0270 |
| DeepLoc Sub. | 0.0120, 0.0100, 0.0110 | 0.0300 |
| EC | 0.0070, 0.0060, 0.0050 | 0.0210 |

### B.2 Confidence Calibration

Reliable confidence estimates are critical for model interpretability and downstream use. We assess calibration using Expected Calibration Error (ECE) for classification and Expected Normalized Calibration Error (ENCE) for regression, each computed over 10 equal-width bins over the confidence or predicted value range. ProtLigand shows better calibration than SaProt in both tasks, with lower ECE (0.043 vs. 0.070) and ENCE (0.048 vs. 0.276). Figure 5 shows that ProtLigand tracks the identity line more closely than SaProt across the confidence bins.

### B.3 Confidence Intervals

To provide a clearer estimate of uncertainty across the three independent fine-tuning runs, we report 95% confidence intervals (CIs) for ProtLigand alongside the mean and standard deviation (Table 4). Confidence intervals are computed using a two-tailed *t*-distribution with *n* = 3.

**Table 4:** Mean ± standard deviation and 95% confidence intervals for ProtLigand across the six benchmark tasks.

| Task | Mean ± SD | 95% CI |
| --- | --- | --- |
| HumanPPI (Acc%) | 90.000 ± 0.5 | [88.76, 91.24] |
| Thermostability ($\rho$) | 0.731 ± 0.02 | [0.680, 0.780] |
| Metal Ion Binding (Acc%) | 77.54 ± 0.18 | [77.09, 77.99] |
| DeepLoc Binary (Acc%) | 94.02 ± 0.21 | [93.50, 94.54] |
| DeepLoc Sub. (Acc%) | 83.88 ± 0.07 | [83.71, 84.05] |
| EC ($F_{\max}$) | 0.887 ± 0.002 | [0.880, 0.890] |

## C ABLATION TESTS

### C.1 Ligand Generator Ablation

To evaluate the importance of the ligand generator component in the ProtLigand architecture, we conduct an ablation experiment by replacing the generator with a random representation drawn from a normal distribution (corresponding in shape to the generated ligand representations).

Table 5 presents the comparison across the six benchmark tasks. Replacing the generator leads to consistent drops in performance across all tasks, particularly in HumanPPI and Thermostability, where ligand context is known to play a key regulatory role. This highlights the contribution of the ligand generator in producing meaningful proxy representations, enabling ProtLigand to capture biochemically relevant signals that guide downstream prediction.

**Table 5:** Ablation study of the ligand generator. We compare the full model to a variant without the ligand generator, replaced by a random ligand representation. Results are reported as mean ± std over 3 independent fine-tuning runs.

| Task | Full Model | Random Ligand |
| --- | --- | --- |
| HumanPPI (Acc%) | 90.00 ± 0.5 | 87.78 ± 0.46 |
| Thermostability ($\rho$) | 0.731 ± 0.02 | 0.716 ± 0.002 |
| Metal Ion Binding (Acc%) | 77.54 ± 0.18 | 74.89 ± 0.25 |
| DeepLoc Binary (Acc%) | 94.02 ± 0.21 | 93.38 ± 0.20 |
| DeepLoc Sub. (Acc%) | 83.88 ± 0.07 | 83.11 ± 0.28 |
| EC ($F_{\max}$) | 0.887 ± 0.002 | 0.878 ± 0.002 |

### C.2 Cross-Attention Ablation

To evaluate the contribution of the cross-attention mechanism in modeling the interaction between protein and ligand representations, we conduct an ablation study in which the cross-attention layer is removed. Instead, we concatenate the protein and ligand representations and project them through a single linear layer to match the protein representation dimension. Specifically, we replace the attention block with: 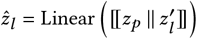 where [[· ∥ ·]] denotes vector concatenation, 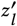 is the ligand representation and *z*_*p*_ is the protein representation (See Section A.1 for details). This configuration removes explicit modeling of interaction patterns between the two modalities, relying instead on a fixed, shallow fusion.

Table 6 presents the comparison across the six benchmark tasks. While the concatenation-based model performs reasonably well, it consistently underperforms the full cross-attention model across all tasks. This performance gap demonstrates that shallow fusion of protein and ligand representations is insufficient for capturing the nuanced, task-specific dependencies between modalities. Nevertheless, the concatenation-based model still outperforms the variant with a random ligand representation (Table 5), indicating that even simpler integration of ligand features is beneficial for downstream performance. The constant improvements with crossattention highlight its role in enriching the protein representations through more informative ligand-contextual interactions.

**Table 6:** Ablation study of the cross-attention mechanism. We compare the full model to a variant where protein and ligand representations are concatenated and linearly projected. Results are mean ± std over 3 independent fine-tuning runs.

| Task | Full Model | Concatenation |
| --- | --- | --- |
| HumanPPI (Acc%) | 90.00 ± 0.5 | 88.33 ± 0.38 |
| Thermostability ($\rho$) | 0.731 ± 0.02 | 0.724 ± 0.004 |
| Metal Ion Binding (Acc%) | 77.54 ± 0.18 | 75.59 ± 0.19 |
| DeepLoc Binary (Acc%) | 94.02 ± 0.21 | 93.55 ± 0.23 |
| DeepLoc Sub. (Acc%) | 83.88 ± 0.07 | 83.33 ± 0.18 |
| EC ($F_{\max}$) | 0.887 ± 0.002 | 0.882 ± 0.002 |

### C.3 Robustness Across Sequence-Identity Bins

We evaluate ProtLigand’s ability to generalize beyond its pre-training distribution by stratifying the HumanPPI test set based on sequence similarity to PDBbind, the corpus used during pre-training. Each HumanPPI test example consists of a protein pair; for each pair, we compute the Needleman–Wunsch sequence identity of both proteins to their respective closest match in the pre-training set. We then average the two identities to obtain a single similarity score for the pair. The full test set is sorted by these pairwise similarity scores and partitioned into five equal-sized bins.

Figure 6 plots the average HumanPPI classification accuracy for each bin against its mean sequence identity. As expected, accuracy increases with similarity, but performance remains strong even in the low-similarity regime: in the lowest bin, where mean identity is just 16.02%, accuracy reaches 83.3%; in the highest bin, where identity approaches 28.5%, performance climbs to 94.4%.

**Figure 6.**
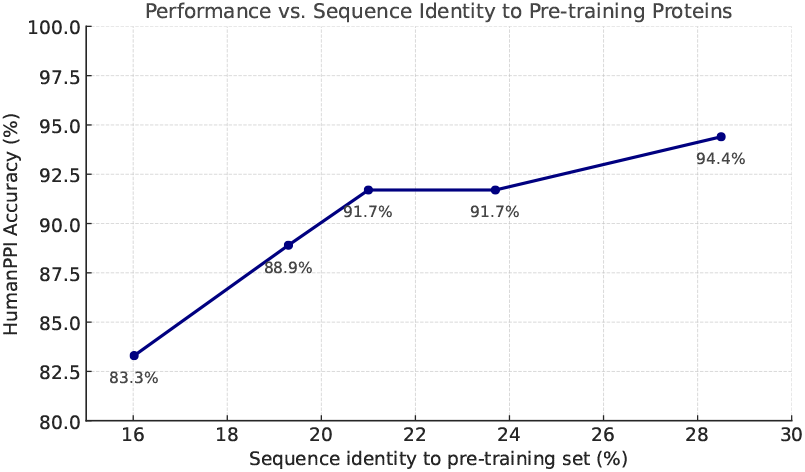
ProtLigand performance on the HumanPPI task as a function of pairwise sequence identity to proteins in the pretraining set. For each test pair, we compute the mean identity of the two proteins to their closest pre-training counterparts. The test set is split into five equal-sized bins by this score. Each point shows the average classification accuracy within a bin, plotted against that bin’s mean identity.

These results confirm that ProtLigand generalizes beyond memorization of homologous sequences. It maintains robust performance even when both proteins in a test pair are dissimilar to any pretraining example, highlighting its potential to operate effectively in real-world contexts such as metabolite-rich or novel ligand datasets, where target proteins may lie well outside the PDBbind manifold.

## D DATASETS OVERVIEW

### D.1 Protein–Ligand Dataset

We provide in Table 7 an overview of the protein–ligand complex distribution by protein type in our pre-training dataset. To better understand the chemical diversity of ligand interactions, we classified each ligand into broad functional categories using the RDKit cheminformatics library, based on their molecular SMILES representations. As shown in Table 9, the dataset spans a broad range of ligand types, including lipids, drug-like molecules, peptides, and nucleotides, reflecting the functional diversity of biologically relevant binding events.

**Table 7:** Total number of protein–ligand complexes for each protein type in the pre-training dataset.

| Protein Type | Protein Amount |
| --- | --- |
| Transferase | 4941 |
| Hydrolase | 4843 |
| Other | 3199 |
| Transcription | 960 |
| Lyase | 848 |
| Transport | 589 |
| Oxidoreductase | 562 |
| Ligase | 434 |
| Isomerase | 320 |
| Chaperone | 297 |
| Membrane | 270 |
| Metal-containing | 112 |
| Viral | 18 |

**Table 8:** Benchmark dataset statistics for the six benchmark tasks.

| Dataset | Category | Evaluation Metric | Train | Valid | Test |
| --- | --- | --- | --- | --- | --- |
| HumanPPI [37] | PPI Prediction | Accuracy | 26319 | 234 | 180 |
| Thermostability [7] | Protein Function Prediction | Spearman’s $\rho$ | 5056 | 639 | 1336 |
| Metal Ion Binding [13] | Protein Function Prediction | Accuracy | 5067 | 662 | 665 |
| DeepLoc (Binary) [1] | Protein Localization Prediction | Accuracy | 5477 | 1336 | 1731 |
| DeepLoc (Subcellular) [1] | Protein Localization Prediction | Accuracy | 8747 | 2191 | 2747 |
| EC [11] | Protein Annotation Prediction | $F_{\max}$ | 13089 | 1465 | 1604 |

**Table 9:** Distribution of ligand types in the pre-training dataset.

| Ligand Type | Count |
| --- | --- |
| Lipid | 6970 |
| Drug-like | 4526 |
| Peptide | 2358 |
| Carbohydrate | 1972 |
| Nucleotide | 1500 |
| Metal-containing | 35 |
| Other | 32 |

### D.2 Benchmark datasets

Table 8 summarizes the benchmark datasets, including task categories, evaluation metrics, and the sizes of the training, validation, and test splits.

### Footnotes

1 The code and datasets used for the statistical testing are freely available in this GitHub repository: https://github.com/kalifadan/ProtLigand

## Notes

https://github.com/kalifadan/ProtLigand

### Competing Interest Statement

The authors have declared no competing interest.

